# Vitamin D receptor and STAT3 cooperate to establish TET2-mediated tolerogenesis

**DOI:** 10.1101/2020.02.28.969634

**Authors:** Francesc Català-Moll, Tianlu Li, Laura Ciudad, Javier Rodríguez-Ubreva, Esteban Ballestar

## Abstract

The active form of vitamin D, 1,25-dihydroxyvitamin D3 (1,25(OH)2D3), induces stable tolerogenesis in dendritic cells (DCs). This process involves the vitamin D receptor (VDR), which translocates to the nucleus, binds its cognate genomic sites, and promotes epigenetic and transcriptional remodeling. In this study, we investigated the interplay between the VDR and other transcription factors to induce DNA methylation changes that might provide phenotypic stability to the tolerogenic phenotype of DCs. Our study reveals the occurrence of vitamin D-specific DNA demethylation and transcriptional activation at VDR binding sites associated with the acquisition of tolerogenesis. Tolerogenic properties in DCs are acquired together with activation of the IL6-JAK-STAT3 pathway. In fact, VDR directly binds the *IL6* gene, and JAK2-mediated STAT3 phosphorylation is specific to vitamin D stimulation. VDR and the phosphorylated form of STAT3 interact with each other and with methylcytosine dioxygenase TET2 following vitamin D treatment. Most importantly, pharmacological inhibition of STAT3 phosphorylation reverts the vitamin-induced tolerogenic properties of DCs. Our results reveal an interplay between VDR and STAT3 leading to the DNA demethylation-dependent induction of tolerogenesis by vitamin D.

## INTRODUCTION

Myeloid cells are not only responsible for innate responses but also participate in initiating adaptive responses. The immunological properties of myeloid cells vary with the environment. In fact, terminal myeloid cell differentiation is highly dependent on the activation of specific signaling pathways in response to extracellular signals, such as inflammatory cytokines, hormones, vitamins and other factors (Álvarez-Errico et al., 2015), which determine the immunogenicity of the resulting myeloid cells. The activation of signaling pathways leads to the activation of specific sets of transcription factors (TFs). Sequence-specific DNA binding of TFs is a pivotal process for establishing gene expression patterns in concert with the epigenetic machinery that determines cell identity and function (Monticelli and Natoli, 2017). Recent evidence has shown that pioneer TFs are associated with DNA demethylation and increased genomic accessibility of their binding genomic regions, facilitating the binding of subsequent TFs (Rasmussen et al., 2019). In this regard, methylcytosine dioxygenase ten-eleven translocation (TET) 2, the most relevant TET enzyme in the myeloid compartment, can interact with a variety of pioneer TFs such as PU.1, C/EBPα, KLF4 and others, which target it to different genomic regions (Costa et al., 2013; Guilhamon et al., 2013; de la Rica et al., 2013; Lio et al., 2016; Sardina et al., 2018; Wang et al., 2015; Xiong et al., 2016). Recently, it has been demonstrated that TET2 mutations, which are frequent in myeloid leukemias, result in enhancer DNA hypermethylation and changes in the subsequent binding of TFs, particularly members of the basic helix-loop-helix (bHLH) TF family (Rasmussen et al., 2019). This suggests that TET2 recruitment by pioneer TFs leads to epigenetic remodeling that facilitates the binding of other TFs (Rasmussen et al., 2019). However, the mechanisms by which this process takes place and the way in which gene expression patterns in primary cell differentiation are acquired and stabilized are poorly understood.

Calcitriol (1,25(OH)2D3), the active form of vitamin D3, is a major modulator of the immune system (Barragan et al., 2015; Carlberg, 2019; Mora et al., 2008). Myeloid cells such as antigen-presenting dendritic cells (DCs) are the target most susceptible to vitamin D in a mixed immune population (Mora et al., 2008). In these cells, calcitriol can generate *in vitro* a stable maturation-resistant tolerogenic phenotype, with a low level of expression of immunogenic molecules such as HLA-DR, CD80 and CD86, and increased interleukin 10 (IL10)/IL12p70 ratios that are maintained even after removal of the compound (Van Halteren et al., 2002). After ligand recognition, vitamin D receptor (VDR) translocates to the nucleus and acts not only as a TF controlling the expression of a set of immune and metabolic genes (Carlberg, 2019; Ferreira et al., 2013), but also as a k-light-chain-enhancer of activated B cell (NF-kB) repressor at different levels (Carlberg, 2019; Fetahu et al., 2014). Several studies have shown the capacity of VDR to interact with a range of TFs including PU.1 and GABPA, and with chromatin remodeling and histone modification enzymes such as BRD7 and KDM6B (Pereira et al., 2011; Seuter et al., 2017, 2018; Wei et al., 2018). There is some evidence that vitamin D may induce DNA methylation alterations (Doig et al., 2013; O’Brien et al., 2018). However, the molecular mechanism that leads to the acquisition of differential methylation patterns and the manner in which autocrine activation of secondary TFs participates in this process remains unexplored.

Vitamin D supplementation is generally used as a preventive agent or a co-adjuvant for diseases with underlying autoimmune or pro-inflammatory states (Bscheider and Butcher, 2016; Dankers et al., 2017). DCs represent an excellent target of vitamin D to dampen autoimmunity and inflammation, not only because these myeloid cells express the whole set of enzymes to generate the active form of vitamin D (Mora et al., 2008) but also for their unique role as initiators of immune responses. However, the role of DCs in vitamin D-mediated immunomodulation is not fully understood. In addition, DCs with tolerogenic function (TolDCs) have become a promising immunotherapeutic tool for reinstating immune tolerance in autoimmune diseases and in allogeneic bone marrow transplantation (Vanherwegen et al., 2019b, 2019a; Willekens et al., 2019). The stability of the tolerogenic phenotype suggests that regulatory mechanisms that allow the maintenance of stable changes of gene expression are involved. In this sense, DNA methylation is a major epigenetic modification closely involved in the acquisition or stabilization of transcriptional states (Luo et al., 2018).

In this study, we studied epigenetic determinants critical for the acquisition of tolerogenic properties during DC differentiation in the presence of vitamin D. We demonstrate an interplay between VDR and the IL6-JAK2-STAT3 pathway in generating a TET2-dependent DNA methylation signature. It involves a direct physical interaction between VDR, STAT3 and TET2 that leads to the acquisition and stabilization of the tolerogenic properties of DCs in the presence of vitamin D.

## RESULTS

### Vitamin D3 induces the acquisition of a specific DNA methylation profile associated with tolerogenesis during DC differentiation

To investigate the participation of VDR and its potential interplay with other TFs in targeting functional DNA methylation changes involved in the acquisition of tolerance, we first differentiated monocytes (MOs) to dendritic cells (DCs) and tolerogenic DCs (TolDCs) *in vitro* for 5 days using GM-CSF and IL-4 in the absence and presence of vitamin D3, respectively (Figure 1A). As previously described (Penna and Adorini, 2000; Piemonti et al., 2000), TolDCs had higher levels of the surface markers CD14 and CD11b and lower levels of HLA-DR, CD1a and CD86 than did DCs (Supplementary Figure 1A). In addition, CD8^+^ T cell proliferation assays in co-culture with both DCs and TolDCs confirmed the immunosuppressive properties of the latter (Supplementary Figure 1B). We also established that vitamin D3 induces the translocation of the VDR to the nucleus under these conditions (Supplementary Figure 1C).

**Figure 1.**
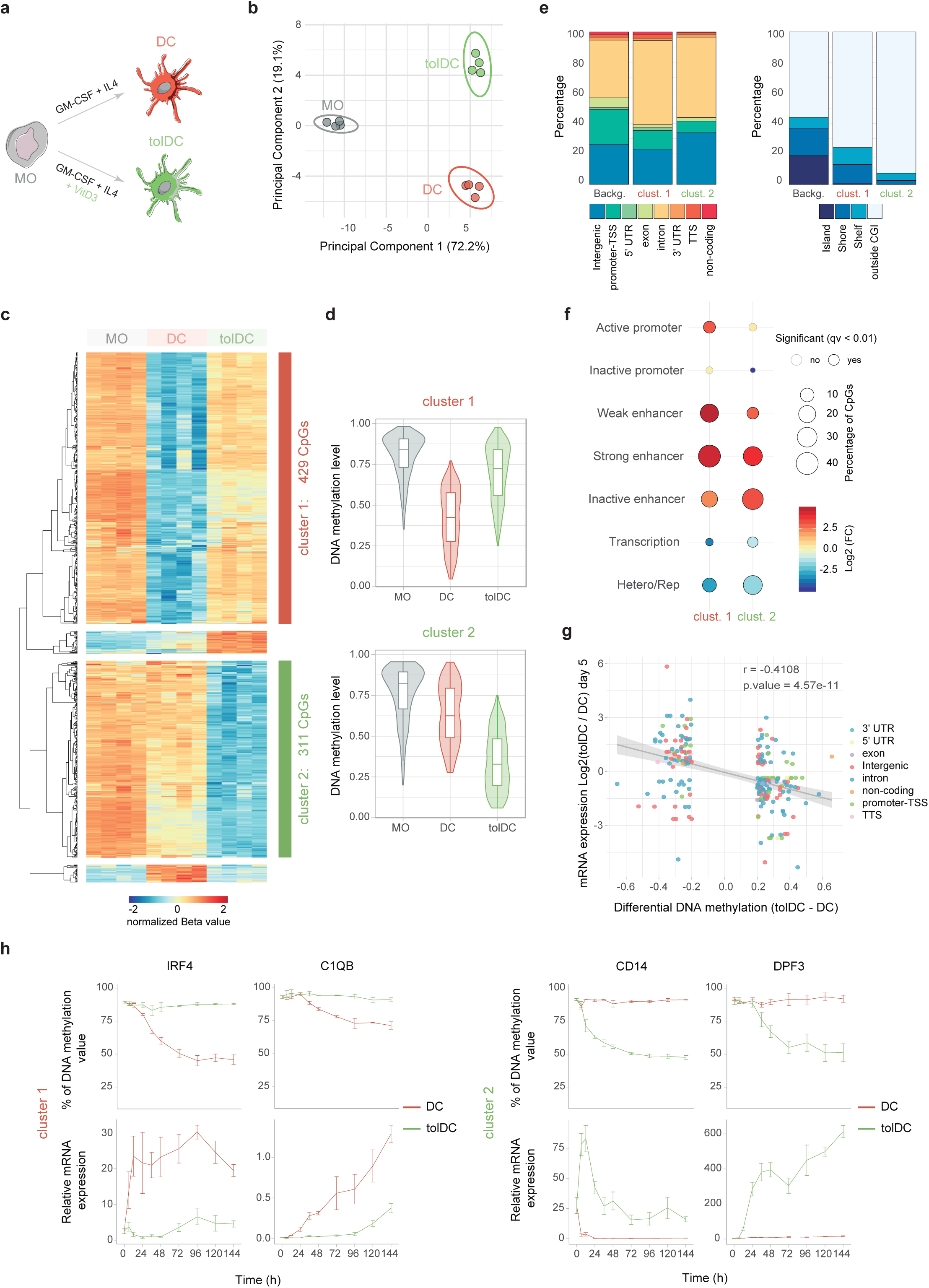
DNA methylation dynamics throughout vitamin D-exposed dendritic-cell differentiation. (**a**) Schematic overview of differentiation model. (**b**) Principal component analysis of differentially methylated CpGs. (**c**) DNA methylation heatmap and cluster analysis of four paired samples of MOs and their derived DCs and TolDCs at day 5 of differentiation. The heatmap includes all CpG-containing probes displaying significant methylation changes (∆β ≥ 0.2 and q < 0.05) only in the TolDC-DC comparison. The color annotation marks the membership of cluster 1 (DC-specific DNA demethylation) or cluster 2 (TolDC-specific DNA demethylation). (**d**) Box and violin plots summarizing the distribution of DNA methylation levels per cell type of cluster 1 (top) and cluster 2 (bottom). (**e**) Location proportions of CpGs of cluster 1 and cluster 2 in the context of CpG islands (CGIs) (right) and gene-related regions (left). (**f**) Bubble chart depicting the enrichment (red) or depletion (blue) of chromatin states on dendritic cells. The circle filling represents the logarithmic-fold change, circle size indicates the percentage of CpGs in the chromatin state, and the circle edge indicates the statistical significance of the enrichment (black: significant; no edge: not significant; q < 0.01). (**g**) Scatter plot showing the correlation between DNA methylation and gene expression (on day 5 of differentiation). Only differentially methylated CpGs are represented. Dot color indicates gene-related associations. (**h**) DNA methylation (top) and mRNA expression (bottom) kinetics of two representative examples of cluster 1 and cluster 2. Statistical tests: two-tailed Fisher’s exact test (f) and Pearson correlation (g).

We performed DNA methylation profiling of MOs and the resulting DCs and TolDCs. Principal component analysis (PCA) showed that most of the variability observed at the DNA methylation level may be explained by events common to the two differentiation processes. However, the second principal component is capable of clustering DCs and TolDCs separately (Figure 1B). Differentiation mainly resulted in DNA demethylation in which common and condition-specific DNA methylation changes were observed, although methylation gains also occurred (Supplementary Figure 2A). Hierarchical clustering of differentially methylated CpGs between DCs and TolDCs (adjusted P < 0.05 and absolute Δß ≥ 0.2) revealed four main groups of CpG sites: a group of CpGs that undergo specific demethylation in DCs (cluster 1, 429 CpGs); a second group that specifically demethylates in TolDCs (cluster 2, 311 CpGs); and two CpG sets with DC- and TolDC-specific gains of DNA methylation (Figure 1C, D). Given the low frequency of CpGs in the hypermethylation clusters, we focused on condition-specific demethylation events (i.e., clusters 1 and 2) in subsequent analyses to investigate how vitamin D treatment leads to changes in the DNA methylome.

Functional gene ontology analysis revealed that CpGs in cluster 1 are associated with immunological categories such as defense and immune response, whereas those in cluster 2 are more highly enriched in cell activation, cell-matrix adhesion, positive regulation of immune system process and wound healing involved in inflammatory response (Supplementary Figure 2B). In both clusters, the majority of changes occurred in introns and intergenic regions with underrepresentation of promoter-transcriptional start sites (TSS). However, while cluster 1 exhibits a marked enrichment of intronic regions with respect to the background, cluster 2 is enriched in intergenic locations (Figure 1E, left). In accordance with this, CpG island annotation showed that CpGs of both clusters are usually located outside of CpG islands, particularly for cluster 2 (Figure 1E, right). Next, we mapped the chromatin states of the CpG sites undergoing demethylation in the two clusters using chromatin segmentation data corresponding to DCs (Pacis et al., 2015) (Figure 1F). We observed an enrichment in the enhancer regions for the two clusters. Moreover, whereas cluster 1 (DC-specific demethylation) is enriched in weak (H3K27ac + H3K4me1 + H3K4me3) and strong enhancers (H3K27ac + H3K4me1), cluster 2 (TolDC-specific demethylation) is more enriched in inactive enhancers in DCs, suggesting that these inactive regions in DCs are activated in TolDCs. To validate this hypothesis, we integrated our DNA methylation dataset with available expression data (Széles et al., 2009). This enabled us to identify a significant inverse relationship between levels of DNA methylation and mRNA expression at 12 h (r = -0.5926, *P* = 4.90e-14) and 5 days of differentiation (r = -0.4108, *P* = 4.57e-11) (Figure 1G and Supplementary Figure 2C). As expected, we were able to confirm that the genes associated with CpGs of cluster 2 located at inactive enhancers of DCs displayed higher expression levels in TolDCs (Supplementary Figure 2D).

To explore the dynamics of the relationship between DC- (cluster 1) and TolDC-specific demethylation (cluster 2), we performed bisulfite pyrosequencing and qRT-PCR of a small selection of genes of a set of samples over time. Bisulfite pyrosequencing showed a high concordance (r = 0.978, *P* < 2.2e-16) with the data obtained from the EPIC arrays (Supplementary Figure 2E). These analyses confirmed not only the specificity of the changes, but also their early occurrence and the concomitant changes between DNA methylation and gene expression (Figure 1H).

### VDR binding is associated with targeted DNA demethylation in TolDCs

We explored the relationship between VDR and the observed DNA methylation changes in DC and TolDC differentiation by performing a ChIP-seq of VDR in our model. The analysis showed that exposure to vitamin D during DC differentiation leads to an increase in VDR genomic binding (Figure 2A,B). Interestingly, motif discovery analysis revealed promiscuity of VDR with respect to its genomic binding preferences, with only 37% of regions having the canonical VDR binding motif (Figure 2C). Functional annotation of associated genes reveals enrichment of immunological- and signaling-related categories (Figure 2D). In fact, a number of genes potentially related to the tolerogenic properties of TolDCs are direct targets of VDR (Supplementary Table 1). Global inspection of VDR genomic occupancy showed that VDR preferentially binds to promoters and introns in comparison with the background (Figure 2E, left). We also observed enrichment of VDR binding in CpG islands, shores and shelves that is compatible with the enrichment noted in promoters (Figure 2E, right). Annotation with the DC chromatin states of VDR peaks confirmed the preference of VDR for binding promoter regions, although it also revealed enrichment in enhancer regions (Figure 2F).

**Figure 2.**
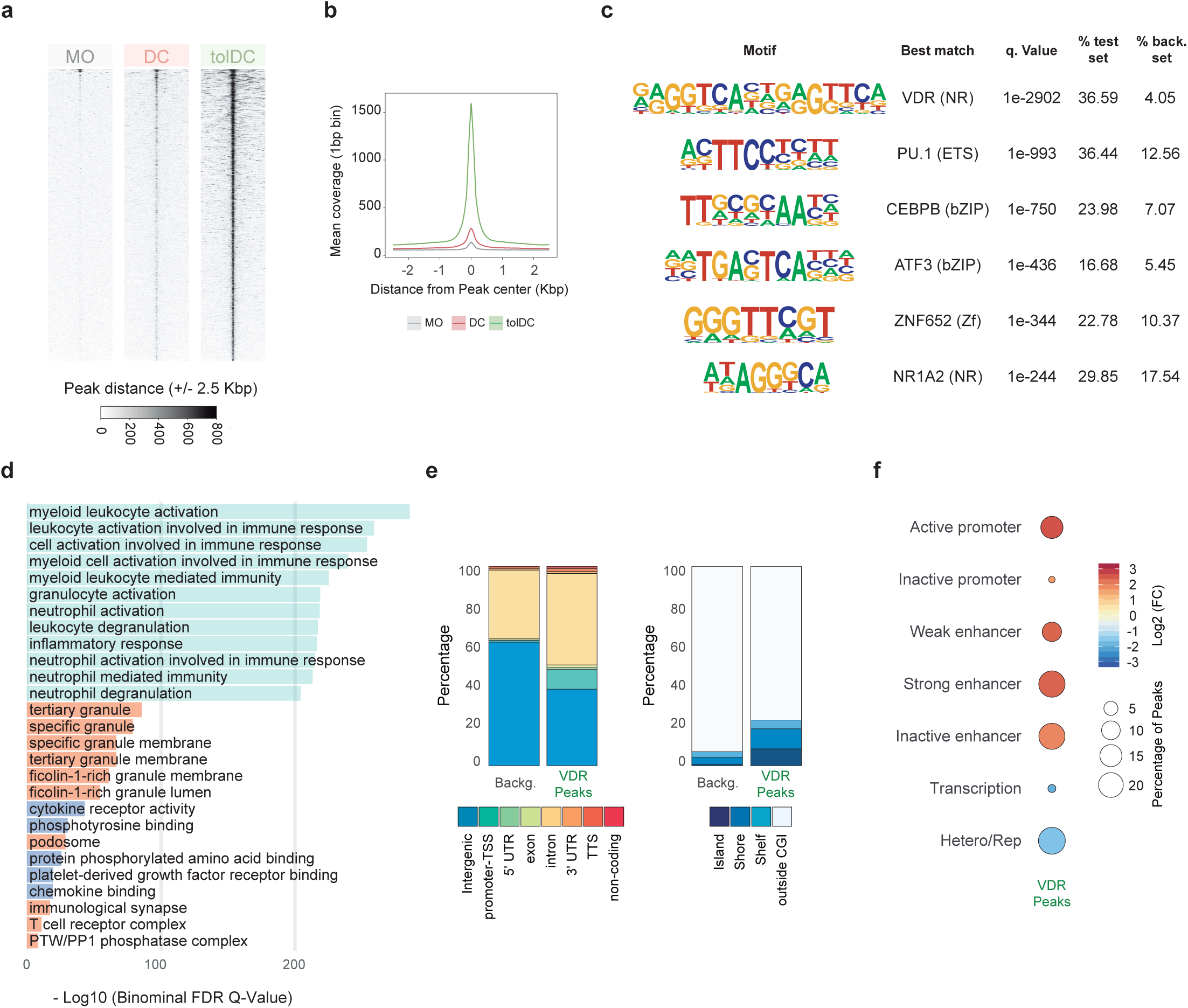
Genomic occupancy of vitamin D receptor. **(a)** Heatmaps showing signal of vitamin D receptor (VDR) ChIP-seq at ± 2.5 Kbp window of significant VDR peaks. **(b)** Composite plots of VDR ChIP-seq distribution ± 2.5 Kbp around CpGs in MO (grey), DC (red) and TolDC (green) for significant VDR peaks. Smooth represents the confidence intervals (CIs). **(c)** Motif discovery analysis using HOMER software, showing q values, percentage of regions with motif and percentage of background sequences with motif. **(d)** Results of gene set enrichment analysis using GREAT software. The plot depicts the top enriched terms for biological processes (green), molecular function (orange) and cellular component (purple) categories, based on adjusted *P* values from the binominal distribution. **(e)** Location proportions of CpGs of cluster 1 and cluster 2 in the context of CpG islands (CGIs) (right) and gene-related regions (left). (**f**) Bubble chart depicting the enrichment (red) or depletion (blue) of chromatin states of dendritic cells. The circle filling represents the logarithmic-fold change, circle size indicates the percentage of CpGs in the chromatin state, and the circle edge indicates the statistical significance of the enrichment (black: significant; no edge: not significant; q < 0.01). Statistical tests: two-tailed Fisher’s exact test (f).

Overlap of DNA methylation and VDR ChIP-seq datasets revealed a specific association between VDR binding and TolDC-specific demethylation (cluster 2) (Figure 3A,B). In fact, we observed that over 40% of CpG sites in cluster 2 had significant VDR binding (Figure 3C). We obtained VDR ChIP-seq signal profiles for several representative genes with CpGs of cluster 2. In general, VDR peaks are close to CpGs that become demethylated in TolDCs (Figure 3D). Bisulfite pyrosequencing revealed that active demethylation of these sites occurred as early as 24 hours following Vitamin D treatment (Figure 3E). In this regard, we have previously shown that loss of methylation in terminal differentiation from MOs is accompanied by a transient increase in 5hmC and involves the participation of TET2 methylcytosine dioxygenase (Garcia-Gomez et al., 2017).

**Figure 3.**
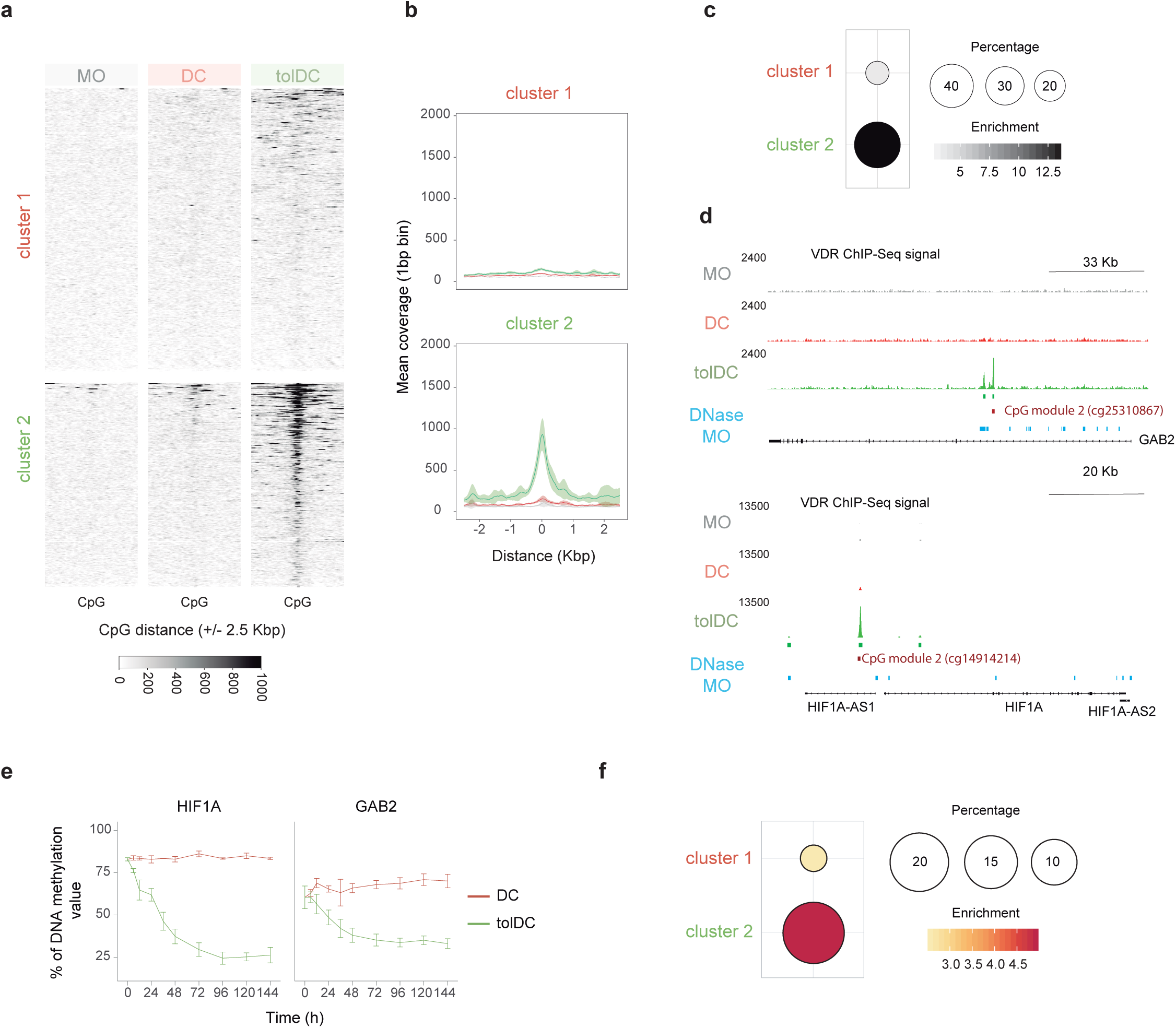
Binding of vitamin D receptor correlates with TolDC-specific DNA demethylation. **(a)** Heatmaps showing signal of vitamin D receptor (VDR) ChIP-seq at ± 2.5 Kbp window of CpGs of cluster 1 (top) and cluster 2 (right). (**b**) Composite plots of VDR ChIP-seq distribution ± 2.5 Kbp around CpGs in MO (grey), DC (red) and TolDC (green) for cluster 1 (top) and cluster 2 (bottom). Smooth represents the confidence intervals (CIs). (**c**) Bubble plot of significant VDR binding enrichment in each cluster of CpGs. Dots are colored according to their enrichment value. Bubble size corresponds to the percentage of CpGs overlapping with significant VDR peaks for each cluster. The presence of a black border indicates significant enrichment (q < 0.01). (**d**) VDR ChIP-seq signal profiles in the vicinity of the representative genes with CpGs of cluster 2. VDR signals are colored by cell type. At the bottom, the significant VDR binding sites are shown in green. CpG position is depicted below in red. Finally, DNase significant peaks in MOs are colored in blue. (**e**) DNA methylation kinetics of the CpGs represented in panel e. **(f)** Bubble plot of significant MO DNase binding enrichment in each cluster of CpGs. Dots are colored by enrichment value. Bubble size corresponds to the percentage of CpGs overlapping with significant DNase peaks for each cluster. The presence of black border indicates significant enrichment (q < 0.01). Statistical tests: two-tailed Fisher’s exact test (c and f).

Hence, we explored the possibility that VDR directly binds to closed chromatin, which may subsequently induce epigenetic remodeling, perhaps through the recruitment of TET2 and/or other epigenetic modifiers. To this end, we performed enrichment analysis comparing DNase-seq datasets from MOs (Feingold et al., 2004) and differentially methylated regions during differentiation to DCs and TolDCs. Our analysis confirmed that over 75% of TolDC-specific demethylated CpG sites overlap with closed regions of MOs (Figure 3F; also see examples in Figure 3D, bottom, and Supplementary Figure 3), which is compatible with a role for VDR as a pioneer TF.

### VDR induces IL-6 upregulation in TolDCs, leading to STAT3 activation and formation of a complex with TET2

Vitamin D3, through its receptor VDR, induces changes in cytokine production and a profound metabolic reprograming (Ferreira et al., 2015). For this reason, we hypothesized that the autocrine activation of secondary signaling pathways during differentiation could lead to the activation of a set of TFs that can cooperate with VDR to mediate changes at the transcriptional and epigenetic levels. To explore this possibility, we adapted a tool initially designed to explore intercellular communication in bulk and single-cell expression data to test autocrine signal activation (Browaeys et al., 2019). With this approach, and using genes associated with both demethylation clusters with significant expression differences (abs(log_2_(FC)) ≥ 1 and adjusted *P* < 0.05) as input, we inferred potential ligands that can explain these differences in expression (Figure 4A). Under these conditions, IL-6 stands out for its role in tumor immune suppression (Park et al., 2017). In fact, the *IL6* gene is significantly more expressed in TolDCs than in DCs (Figure 4B) and this upregulation could play a major role in the expression of target genes (Figure 4C).

**Figure 4.**
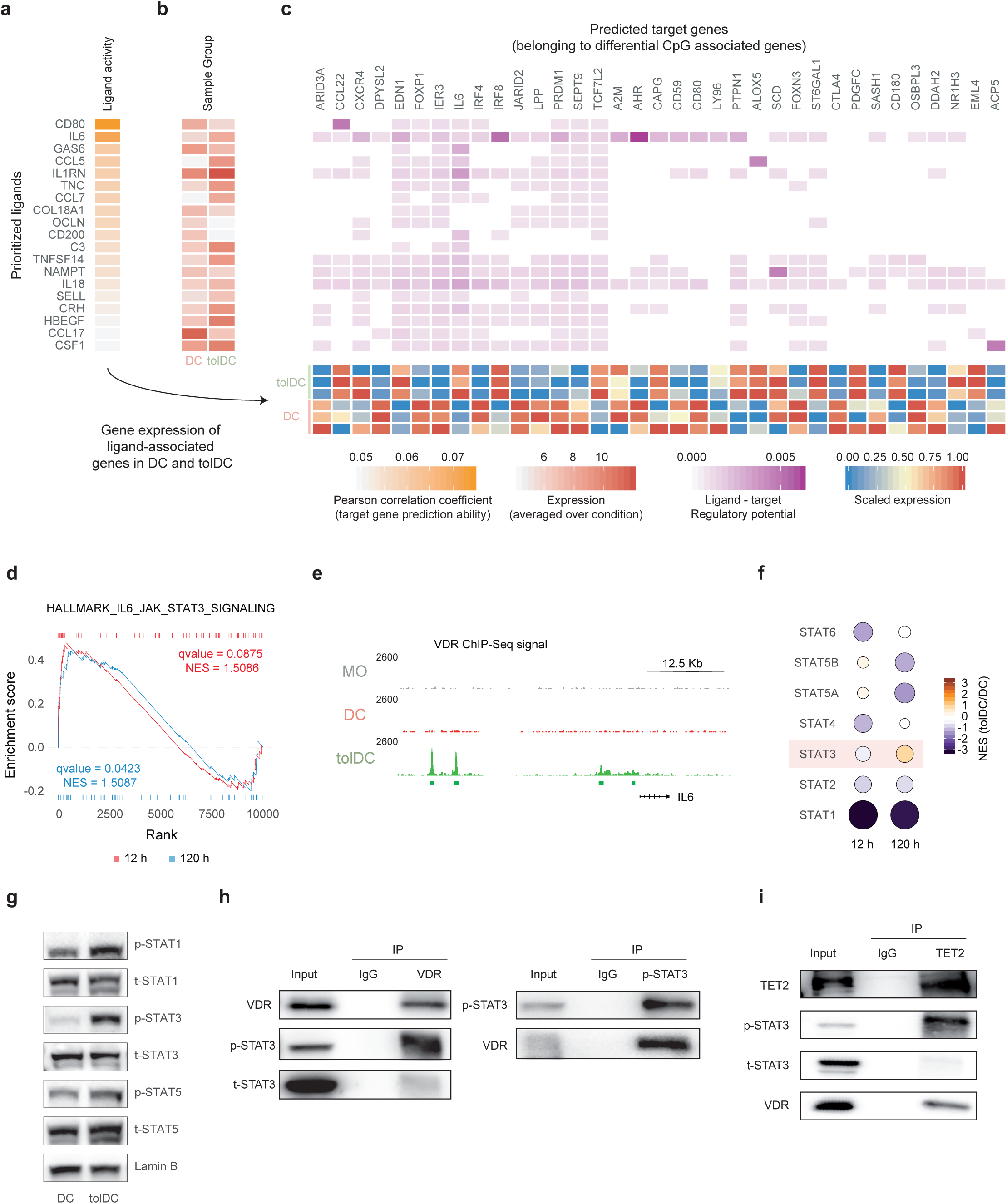
Vitamin D-dependent autocrine activation of the IL-6/JAK2/STAT3 pathway. (**a**) Heatmap showing ligand activity based on the Pearson correlation with its target genes. (**b**) Heatmap displaying average gene expression of ligands for DCs and TolDCs on day 5. (**c**) Heatmap showing the regulatory potential for each ligand on the target genes (upper panel) and its expression levels in each sample (bottom). (**d**) Gene set enrichment analysis of differentially expressed genes (abs(log_2_(FC)) ≥ 1 and q < 0.05) at 12 h (red) and 120 h (blue). Results for the IL-6/JAK/STAT signaling pathway are shown. (**e**) VDR ChIP-seq signal profiles in the vicinity of the IL-6 gene. VDR signals are colored by cell type. The significant VDR binding sites are shown below in green. (**f**) Schematic representation of the IL-6/JAK2/STAT3 pathway. (f) Bubble chart depicting the TF activity predicted from mRNA expression with DoRothEA v2.0. The circle filling represents the normalized enrichment score (NES) (blue: more activity in DCs, red: more activity in TLs). Bubble size corresponds to the logarithm of adjusted *P* values. (**g**). WB showing the phosphorylated and total protein levels of STAT1, STAT3 and STAT5 on day 3 of differentiation. (**h, i**) Co-immunoprecipitation assays were performed in MOs differentiated to TolDC for 3 days. Protein extracts were immunoprecipitated using anti-VDR, anti phospo-STAT3 and anti-TET2 antibodies, in which IgG was used as a negative control and total protein extract was used as input.

As a validation, gene set enrichment analysis (GSEA) of differentially expressed genes also showed enrichment of the IL-6/JAK/STAT3 signaling pathway (Figure 4D). In fact, VDR binds in several regions upstream of the *IL6* gene TSS, suggesting that VDR directly regulates its expression (Figure 4E). To establish whether autocrine IL-6 signaling induction activates STAT3, we tested its activity using DoRothEA (Discriminant Regulon Expression Analysis), a manually curated human regulon for estimating single-sample TF activities through the expression of their target genes (Garcia-Alonso et al., 2019). We observed a specific increase in STAT3 activity in TolDCs after 5 days (Figure 4F). We then tested the phosphorylation levels of STAT3 in parallel with other STATs (STAT1 and STAT5) 3 days after differentiation in the absence or presence of vitamin D3. STAT3 showed specific phosphorylation in TolDCs but not in DCs. This behavior differed from that observed for STAT1 and STAT5, which were phosphorylated in both conditions (Figure 3G). Thus, our results indicate that vitamin D induces the autocrine activation of IL-6 signaling with subsequent STAT3 activation. Interestingly, a similar involvement of the IL-6/JAK/STAT3 pathway was found in *in vitro* differentiated myeloid derived suppressor cells, obtained by adding prostaglandin E2 (PGE2) during differentiation to DCs (Rodríguez-Ubreva et al., 2017) (Supplementary Figure 4A-C). These alternative tolerogenic DCs also showed increased nuclear levels of the phosphorylated form of STAT3 (Supplementary Figure 4D).

To explore the possibility that the observed interplay between VDR and STAT3 involves a physical interaction, we performed co-immunoprecipitation experiments in TolDCs. Our analysis revealed a specific interaction between VDR and phospho-STAT3 (Figure 4H). We also observed that both VDR and phospho-STAT3 interact with TET2 (Figure 4I), which suggest that these two TFs play a role in the targeting of TET2-mediated demethylation to their cognate sites. In all, our results support the notion that both factors act in a complex to induce the specific demethylation observed in TolDCs.

### Inhibition of STAT3 activation affects the acquisition of vitamin D-dependent tolerogenesis

We investigated the consequences of inhibiting the IL6/JAK/STAT3 pathway by using TG101348, a pharmacological inhibitor of JAK2, during DC and vitamin D-dependent TolDC differentiation. Following TG101348 treatment, we confirmed the inhibition of STAT3 phosphorylation by western blot (Figure 5A). TG101348 treatment also resulted in a sharp decrease in the production of IL-10 (Figure 5B), an archetypical anti-inflammatory cytokine which is also a *bona fide* target for STAT3 (Schaefer et al., 2009; Ziegler-Heitbrock et al., 2003). We also tested the effects of JAK2 inhibition on surface markers. JAK2 inhibition resulted in an increase of CD14 and CD86 protein levels and downregulation of CD1a and CD11b (Figure 5C). In parallel, we investigated the effects of JAK2 inhibition on the DNA methylation and expression levels of TolDC-specific demethylated genes. We did not observe any clear reversion of DNA demethylation (Figure 5D), but we did note alterations at the transcriptional level. Changes were observed not only in genes in cluster 2 (TolDC-specific) but also in those of cluster 1 (Figure 5E). This is likely to be the result of the inhibition of phosphorylation of STAT1 and STAT5, which might also be involved in activating these and other DC and TolDC genes.

**Figure 5.**
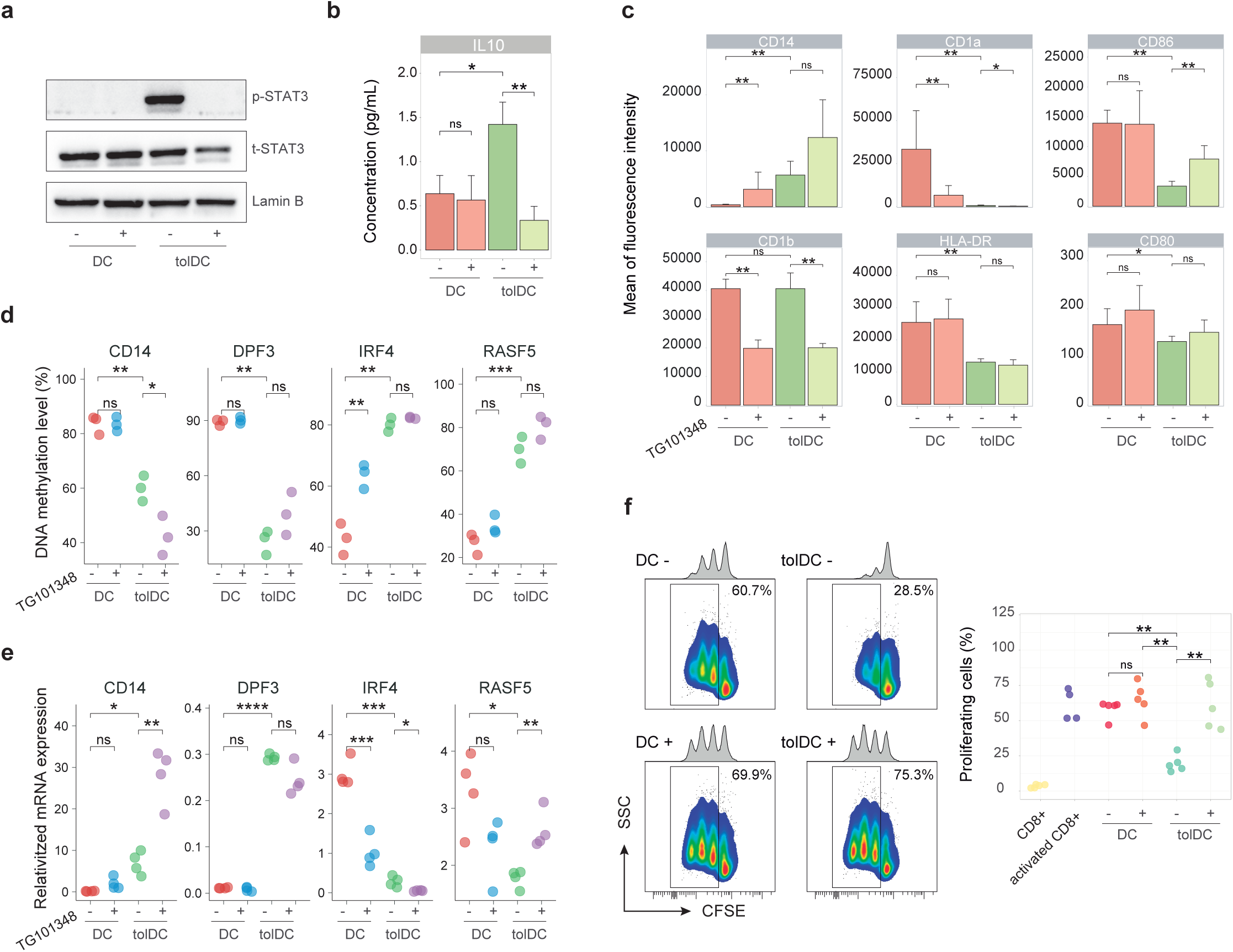
Phosphorylation inhibition of STAT3 reverts immunosuppressive proprieties of vitamin D exposed dendritic cells. (**a**) Effects of STAT3 at the protein phosphorylation level after pharmacological inhibition of JAK2 with TG101348. (**b**) Effect of JAK2 inhibition on IL-10 released by DCs and TolDCs (n = 3). Protein levels were measured by ELISA. (**c**) Bar plots showing the impact of JAK2 inhibition on membrane receptor expression (n = 4). Protein levels were measured with flow cytometry. (**d**) Bisulfite pyrosequencing of four example CpGs displaying the consequence of JAK2 inhibition (n = 3). (**e**) mRNA expression of four example genes from cluster 1 and cluster 2, showing the effect of JAK2 inhibition (n = 4). Expression was relativized with respect to RPL38 gene expression. (**f**) Representative example and dot plot showing the effect on CD8^+^ cell proliferation of DCs and TolDC generated from MO in presence or absence of TG101348 (n = 5). Statistical tests: Two-tailed unpaired Wilcoxon’s test (b-f).

Most importantly, JAK2 inhibition resulted in the loss of the tolerogenic properties of DC differentiated in the presence of vitamin D. In fact, TolDC differentiated in the presence of TG101348 stimulated CD8^+^ cell proliferation to a similar extent to DCs, in contrast with TolDC in the absence of TG101348, reinforcing the idea that the activities of VDR and the JAK2/STAT3 pathway are coordinated in the acquisition of tolerogenic properties (Figure 5F).

## DISCUSSION

In this study, we demonstrate that vitamin D is able to induce tolerogenesis in DCs through a mechanism that involves VDR-specific demethylation and activation of key immune genes in a manner that is coordinated with STAT3 activation. VDR is able not only to orchestrate a direct response on key immune targets but also to mediate a IL6/JAK/STAT-mediated response. This involves the direct binding and activation of the IL6 gene and the recruitment of TET2 by VDR and the phosphorylated form of STAT3 to demethylate and subsequently activate target genes. The essential role of the JAK2-STAT3 pathway in the acquisition of tolerogenesis is demonstrated by the functional impact of the pharmacological inhibition of this pathway.

Our results show the direct role of VDR in guiding TET2-mediated DNA demethylation to specific genomic sites during TolDC differentiation. We have shown that, in the presence of vitamin D3, VDR can bind closed chromatin in MOs and form a complex with TET2, thereby promoting TolDC-specific demethylation. These two findings are compatible with a role of VDR as a pioneer TF (Mayran and Drouin, 2018). This finding is at odds with the findings of a previous study of the human monocytic THP-1 cells, in which the authors found that VDR only binds to open regions before vitamin D3 exposure (Abdelrasoul et al., 2015). However, it is easy to envision that the accessibility profile of immortalized cells could be substantially different from that of normal cells. In addition, the interactions of VDR with different chromatin remodelers (Nurminen et al., 2018, 2019; Pereira et al., 2011; Wei et al., 2018) support the potential role of VDR as a pioneer factor.

A recurrent question in the DNA methylation field is whether DNA methylation is causally involved in shaping gene expression profiles, or if it passively reflects transcriptional states (Schübeler, 2015). Data support both possibilities and some DNA methylation changes appear to be more likely to cause subsequent changes than others. In our study, we present evidence that TET2-mediated demethylation acts as a mechanism facilitating subsequent participation of other TFs, in this case STAT3. In fact, the absence of interference with DNA demethylation, while activation is impeded following pharmacological inhibition of STAT3 phosphorylation, suggests that VDR-dependent demethylation is necessary and precedes STAT3-mediated gene activation. This proposed mechanism was consistent with the alterations in TF activity reported in TET2 knockout mice (Rasmussen et al., 2019). TET2-associated functions may ensure the binding of some TFs, thereby contributing to enhancer-dependent activity and gene expression.

Our study identifies a crucial role for the IL6/JAK/STAT3 pathway in the acquisition of tolerogenesis in innate immunity. The involvement of STAT3 is also relevant in the context of myeloid-derived suppressor cells (MDSCs) and tumor-associated macrophages (TAMs), which are also characterized by their tolerogenic properties (Corzo et al., 2009; Kumar et al., 2016). This pathway plays a role in tumor microenvironment-induced dysregulated immune responses (Park et al., 2017), where IL6/JAK/STAT3 activation is associated with adverse host inflammatory responses and reduced survival. It has been suggested that this pathway is an important immunosuppressive event blocking effective T-cell immune responses in glioma (Yao et al., 2016). Our data shed light on how this pathway is directly activated in the face of vitamin D-mediated immunosuppression, in the context of DCs, which are essential in T-cell-mediated immunity. VDR directly targets the activation of the *IL6* gene, and vitamin D results in the specific phosphorylation of STAT3. We show that the pharmacological impairment of STAT3 phosphorylation, by inhibiting JAK2, directly results in the loss of the tolerogenic properties of TolDCs, which facilitate T-cell proliferation, demonstrating the essential role of this pathway. Our results raise the possibility that tolerogenic properties can be reverted, not only in the context of vitamin D, but also in others.

## EXPERIMENTAL PROCEDURES

### Differentiation of TolDCs and DCs from peripheral blood mononuclear cells

For *in vitro* differentiation experiments, we obtained buffy coats from anonymous donors through the Catalan Blood and Tissue Bank (CBTB). The CBTB follows the principles of the World Medical Association (WMA) Declaration of Helsinki. Before providing the first blood sample, all donors received detailed oral and written information, and signed a consent form at the CBTB. PBMCs were isolated by Ficoll-Paque gradient centrifugation. MOs were isolated from PBMCs using positive selection with MACS CD14 microbeads (Miltenyi Biotec). Cells were resuspended in RPMI Medium 1640 + GlutaMAXTM-1 (Gibco, Life Technologies) containing 10% fetal bovine serum, 100 units/mL penicillin, and 100 μg/mL streptomycin. For TolDC differentiation, the medium was supplemented with 10 ng/mL human IL-4, 10 ng/mL GM-CSF (PeproTech), and 10 nM of vitamin D3 or calcitriol (Sigma Aldrich). For DCs, the medium did not contain vitamin D3. In some cases, differentiation was performed in the presence of a JAK2 inhibitor (TG101348, STEMCELL) at 500 nM.

### CD8^+^ cell proliferation assay

Allogenic CD8^+^ T-cells were isolated by negative selection using Dynabeads Untouched Human CD8 T Cells Kit (Invitrogen), labeled with carboxyfluorescein succinimidyl ester (CFSE) and seeded in 96-well plates at 200,000 cells/well, with TolDCs or DCs at different ratios (TolDC/DC:CD8+ T-cell ratios: 1:2, 1:4, and 1:6). CD8^+^ cells were then stimulated with anti-CD3/CD28 Dynabeads 5 μL/mL (Invitrogen) and cultured for 3 days. CD8^+^ T-cell proliferation was analyzed by FACS and determined by considering the proliferating CD8^+^ T-cells those where CFSE staining had decreased compared to unstimulated CD8^+^ T-cells.

### Cytokine measurements

For *in vitro* experiments, the concentration of cytokines was measured from the cell culture supernatants using an enzyme-linked immunosorbent assay (ELISA), according to the manufacturer’s instructions (BioLegend, San Diego, CA, USA).

### Genomic DNA extraction

DNA was extracted with a Maxwell RSC Cultured Cells DNA kit (Promega).

### Bisulfite pyrosequencing

500 ng of genomic DNA was converted with an EZ DNA Methylation-Gold kit (Zymo Research), following the manufacturer’s instructions. Bisulfite-treated DNA was PCR-amplified using primers (see Table S2) designed with PyroMark Assay Design 2.0 software (Qiagen). Finally, PCR amplicons were pyrosequenced with the PyroMark Q24 system and analyzed with PyroMark CpG software (Qiagen).

### Co-immunoprecipitation (Co-IP)

Co-IP assays were performed using TLs differentiated from CD14+ monocytes for 3 days. Cell extracts were prepared in lysis buffer [50 mM Tris–HCl, pH 7.5, 1 mM EDTA, 150 mM NaCl, 1% Triton-X-100, protease inhibitor cocktail (cOmplete™, Merck)] with corresponding units of Benzonase (Sigma) and incubated at 4°C for 4 h. 50 μl of supernatant was saved as input and diluted with 2× Laemmli sample buffer (4% SDS, 20% glycerol, 120 mM Tris–HCl, pH 6.8). Supernatant was first incubated with PureProteome™ Protein A/G agarose suspension (Merck Millipore) for 1 h to remove background signal. Samples were then incubated overnight at 4°C with corresponding antibodies against VDR (12550, Cell Signaling), TET2 (ab124297, Abcam), STAT3 (79D7, Cell Signaling) and pSTAT3 (D3A7, Cell Signaling) according to the specifications of each antibody. Negative controls were incubated with rabbit (12-370, Merck Millipore) and mouse (12-371, Merck Millipore) IgGs. Subsequently, samples were incubated with magnetic beads at 4°C for 2 h, and beads were then washed three times with lysis buffer. For sample elution, 100 μl of 1× Laemmli sample buffer was added to the beads. Samples and inputs were denatured at 95°C in the presence of 1% β-mercaptoethanol.

### DNA methylation profiling

Infinium MethylationEPIC BeadChip (Illumina, Inc., San Diego, CA, USA) arrays were used to analyze DNA methylation. This platform allows > 850,000 methylation sites per sample to be interrogated at single-nucleotide resolution, covering 99% of the reference sequence (RefSeq) genes. The samples were bisulfite-converted using EZ DNA Methylation-Gold™ Kit (Zymo Research, Irvine, CA, USA) and were hybridized in the array following the manufacturer’s instructions. Image processing and intensity data extraction software and procedures were as previously described. Each methylation data point was obtained from a combination of the Cy3 and Cy5 fluorescent intensities from the methylated and unmethylated alleles. Background intensity computed from a set of negative controls was subtracted from each data point. For representation and further analysis, we used beta and M values. The beta value is the ratio of the methylated probe intensity to the overall intensity (the sum of the methylated and unmethylated probe intensities). It can take a value between 0 and 1, and was used to derive heatmaps and to compare DNA methylation percentages from bisulfite-pyrosequencing experiments. The M value is calculated as the log_2_ ratio of the intensities of the methylated versus unmethylated probes. For the purpose of statistical analysis, M values are more suitable because they are normally distributed.

Raw methylation data were preprocessed with the minfi package (Aryee et al., 2014). Data quality was assessed using the minfi and RnBeads packages (Aryee et al., 2014; Assenov et al., 2014; Müller et al., 2019). After Snoob normalization, data were analyzed using an eBayes moderate t-test available in the limma package. Several criteria have been proposed as representing significant differences in methylated CpGs, but in this study we considered a probe to be differentially methylated if it had a methylation differential of 20% and if it was significant (q < 0.05).

### ChIP-seq analysis

Chromatin immunoprecipitation was performed using the iDeal ChIP-seq kit for Transcription Factors (Diagenode), according to the manufacturer’s instructions. Briefly, cells on day 3 of differentiation were cross-linked with 1% formaldehyde for 15 min and glycine was added to quench the reaction (final concentration 125 mM, incubated for 5 min at room temperature). Cells were washed once with cold PBS, scraped off the plates, and pelleted. To obtain a soluble chromatin extract, cells were resuspended in 1 ml LB1 (50 mM HEPES, 140 mM NaCl, 1 mM EDTA, 10% glycerol, 0.5% NP-40, 0.25% Triton X-100 and 1× complete protease inhibitor) and incubated while rotating at 4°C for 10 min. Samples were centrifuged, resuspended in 1 ml LB2 (10 mM Tris-HCl pH 8.0, 200 mM NaCl, 1 mM EDTA, 0.5 mM EGTA and 1× complete protease inhibitor) and incubated while rotating at 4°C for 10 min. Finally, samples were centrifuged, resuspended in 1 ml LB3 (10 mM Tris-HCl pH 8.0, 100 mM NaCl, 1 mM EDTA, 0.5 mM EGTA, 0.1% sodium deoxycholate, 0.5% *N*-lauroylsarcosine, 1% Triton X-100 and 1× complete protease inhibitor). Chromatin extracts were sonicated for 12.5 min using a Covaris M220 focused ultrasonicator at a peak power of 75, and a duty factor of 10 and 200 cycles per burst. The lysates were incubated with anti-VDR antibody (12550, Cell Signaling) bound to 30 μl protein A or protein G Dynabeads and incubated overnight at 4°C, keeping 5% as input DNA. Magnetic beads were sequentially washed with low-salt buffer (150 mM NaCl, 0.1% SDS, 1% Triton X-100, 1 mM EDTA and 50 mM Tris-HCl), high-salt buffer (500 mM NaCl, 0.1% SDS, 1% Triton X-100, 1 mM EDTA and 50 mM Tris-HCl), LiCl buffer (150 mM LiCl, 0.5% sodium deoxycholate, 0.1% SDS, 1% Nonidet P-40, 1 mM EDTA and 50 mM Tris-HCl) and TE buffer (1 mM EDTA and 10 mM Tris-HCl). For ChIP-seq, beads were resuspended in elution buffer (1% SDS, 50 mM Tris-HCl pH 8.0, 10 mM EDTA and 200 mM NaCl) and incubated for 30 min at 65°C. After centrifugation, the eluate was reverse-cross-linked overnight at 65°C. The eluate was then treated with RNaseA for 1 h at 37°C and with Proteinase K (Roche) for 1 h at 55°C and the DNA was recovered using a Qiagen PCR purification kit.

Sequencing reads from ChIP-seq experiments were mapped to the hg19 assembly of human reference genome using Bowtie2 Aligner v2.2.6 (Langmead et al., 2009). After removing reads with MAPQ < 30 with Sequence Alignment/Map (SAMtools) v1.2 (Li et al., 2009), PCR duplicates were eliminated using the Picard function available in MarkDuplicates software v1.126 (Broad Institute, 2018). Peak calling was determined using SPP (with parameters –npeak=300000 –savr –savp -rf). The irreproducible discovery rate (IDR) was used to filter peaks (IDR < 0.05). To visualize individual ChIP-seq data on Integrative Genomics Viewer (IGV), we converted bam output files to normalized bigwig format using the bamCoverage function in deepTools (v2.0).

### Data analysis

Hierarchical clustering was carried out based on Pearson correlation distances and average linkage criteria. For low-dimensional analysis, we used principal component analysis (PCA). Transcription-factor motifs were enriched for each set using HOMER software v4.10.3. Specifically, we used the findMotifsGenome.pl algorithm (with parameters -size 200 -cpg) to search for significant enrichment against a background sequence adjusted to have similar CpG and GC contents. Genomic regions for genetic context location were annotated using the annotatePeaks.pl algorithm in the HOMER v4.10.3 software application. To determine the location relative to a CpG island (CGI), we used ‘hg19_cpgs’ annotation in the annotatr v1.8 R package. GREAT software (McLean et al., 2010) was used to enrich downstream pathways and gene ontologies. We used the single nearest gene option to identify associations between genomic regions and genes. Chromatin state analysis for DCs were assessed using the EpiAnnotator R package (Pageaud et al., 2018). Inference of TF activities from expression values were calculated using DoRothEA (Garcia-Alonso et al., 2019). We used the nichenetr package (Browaeys et al., 2019) to predict ligand activity.

### Statistical analysis

All statistical analyses were done in R v3.5.1. Data distributions were tested for normality. Normally distributed data were tested using two-tailed unpaired Student’s t-tests; non-normal data were analyzed with the appropriate non-parametric statistical test. Levels of significance are indicated as: *, *P* < 0.05; **, *P* < 0.01; ***, *P* < 0.001; ****, *P* < 0.0001. Non-significance (*P* ≥ 0.05) is indicated as ‘ns’.

## DATA ACCESS

DNA methylation and ChIP-seq data for this publication have been deposited in the NCBI Gene Expression Omnibus and are accessible through GEO Series accession numbers GSE145483 and GSE145584.

## Author contributions

F.C.-M. designed and performed the experiments and bioinformatic analysis; F.C.-M. and T.L. performed the co-immunoprecipitation experiments; L.C. gave technical support; J.R.-U and E.B. supervised the study; E.B. conceived the study; F.C.-M. and E.B. wrote the manuscript; all authors participated in discussions and interpreting the results.

